# Natural selection on gene-specific codon usage bias is common across eukaryotes

**DOI:** 10.1101/292938

**Authors:** Zhen Peng, Hani Zaher, Yehuda Ben-Shahar

## Abstract

Although the actual molecular evolutionary forces that shape differences in codon usage across species remain poorly understood, majority of synonymous mutations are assumed to be functionally neutral because they do not affect protein sequences. However, empirical studies suggest that some synonymous mutations can have phenotypic consequences. Here we show that in contrast to the current dogma, natural selection on gene-specific codon usage bias is common across Eukaryota. Furthermore, by using bioinformatic and experimental approaches, we demonstrate that specific combinations of rare codons contribute to the spatial and sex-related regulation of some protein-coding genes in *Drosophila melanogaster.* Together, these data indicate that natural selection can shape gene-specific codon usage bias, which therefore, represents an overlooked genomic feature that is likely to play an important role in the spatial and temporal regulation of gene functions. Hence, the broadly accepted dogma that synonymous mutations are in general functionally neutral should be reconsidered.

## INTRODUCTION

Most amino acids are encoded by at least two interchangeable synonymous DNA codons (Nirenberg and Leder, 1964; Crick, 1966; Söll et al., 1966). Although synonymous codons do not impact the amino-acid sequences of proteins, analyses of genomic DNA indicate that protein-coding genes often exhibit codon usage bias, which is defined as usage frequencies of specific synonymous codons that deviate from expected random usage frequencies (Ikemura, 1981; Sharp et al., 1988; Plotkin and Kudla, 2011; Doherty and McInerney, 2013). While the specific patterns of codon usage bias often vary across species, this phenomenon seems to be a universal feature of most protein-coding DNA sequences (Sharp et al., 1988; Kanaya et al., 1999; Wang and Hickey, 2007; Quax et al., 2015; Pouyet et al., 2016). Nevertheless, the actual molecular forces that shape gene-specific codon usage bias at the evolutionary timescale remain poorly understood.

Previous theoretical models derived from population genetics suggested that observed codon usage frequencies are the result of an interplay between three forces: 1) Mutational bias, which refers to the asymmetric occurrence of mutations; 2) Genetic drift, whereby specific synonymous codons are randomly fixed; 3) Natural selection for favored specific synonymous codon usage patterns associated with specific gene functions. This theoretical framework was mathematically formalized as the “selection-mutation-drift model” (Bulmer, 1991; Yang and Nielsen, 2008). Although the “selection-mutation-drift model” does not assign different mathematical weights to these three forces, it is generally assumed that mutational bias is likely the dominant force in shaping codon usage bias across diverse species (Stenico, Lloyd, and Sharp, 1994; Chen et al., 2004; Fu, 2010; Sharp, Emery, and Zeng, 2010; Rao et al., 2011; Doherty and McInerney, 2013; Yang et al., 2015). In contrast, whether genetic drift and natural selection play equally important roles in driving gene-specific codon usage patterns remains controversial (Akashi, 1995; Rao et al., 2011; Doherty and McInerney, 2013; Kessler and Dean, 2014). Specifically, since synonymous mutations do not affect protein sequences, and therefore are usually assumed to have no impact on their structure or biochemical function, the effect of natural selection on gene-specific codon usage bias is often dismissed as weak (King and Jukes, 1969; Kimura, 1968; Kimura and Ohta, 1974; Echave, Spielman, and Wilke, 2016).

Nonetheless, a series of biochemical and molecular studies of specific protein-coding genes seem to present mechanistic challenges to the broadly accepted dogma that synonymous mutations are generally functionally neutral. First, gene-specific synonymous codon usage patterns have been shown to be important for the function of some protein-coding genes by affecting features such as translation efficacy (Ikemura, 1981; Akashi, 1994; Ko et al., 2005), mRNA secondary structures (Hasegawa, Yasunaga, and Miyata, 1979), mRNA splicing patterns (Pagani, Raponi, and Baralle, 2005; Parmley and Hurst, 2007), miRNA binding accessibility (Gu et al., 2012), protein folding (Zhang, Hubalewska, and Ignatova, 2009; Zhou et al., 2013; Fu et al., 2016), and post-translational modifications (Kimchi-Sarfaty et al., 2007; Zhou et al., 2013; Quax et al., 2015; Fu et al., 2016). Second, the temporal regulation of protein synthesis in several viruses in the *Herpesviridae* group has been shown to depend on viral-specific codon usage patterns that are different from those of the host cell (Shin et al., 2015). Third, high protein levels of some housekeeping genes depend on the biased usage of common synonymous codons (Stenico, Lloyd, and Sharp, 1994; Ma et al., 2014). Some of these anecdotal examples could be explained by the “translational selection theory”, which stipulates that “optimal” protein translation rate and accuracy partly depend on the availability and affinity of cognate tRNAs, and therefore, necessitates the evolution of codon usage patterns that match the relative abundance of specific tRNAs in the relevant cellular pools (Ikemura, 1981; Shields and Sharp, 1987; Slimko and Lester, 2003; dos Reis, Savva, and Wernisch, 2004). Together, these studies suggest that gene-specific codon usage biases might be playing a general, yet underappreciated, role in the spatial and temporal regulation of gene activity, and therefore should be affected by natural selection.

Here we combine theoretical and experimental approaches to test the hypothesis that natural selection has played a significant and important role in shaping gene-specific codon usage patterns. To test this hypothesis, we developed a statistical approach for identifying signatures of natural selection on gene-specific codon usage patterns. Using this approach, we demonstrate that a considerable proportion of protein-coding genes from diverse eukaryotic species display signatures of natural selection on biased gene-specific codon usage patterns. Additionally, we use the well-annotated genome of *Drosophila melanogaster* to identify genome-wide signatures of selection on gene-specific codon usage patterns and demonstrate that a specific combination of unique rare synonymous codons are associated with specific biological functions, and specifically enhance protein expression in the male reproductive system. Together, our results indicate that in contrast to the broadly accepted dogma, natural selection plays a significant role in determining gene-specific codon usage biases in diverse eukaryotic species, and that gene-specific codon usage biases are important for the spatial and temporal regulation of some protein-coding genes. Together, these data indicate that the broadly accepted assumption that synonymous mutations are in general functionally and evolutionarily neutral should be empirically evaluated on gene-specific bases.

## RESULTS

### Signatures of natural selection on gene-specific codon usage are common across eukaryotes

Testing the hypothesis that gene-specific codon usage bias plays a general role in regulating gene function requires a broadly applicable method for statistically identifying signatures of selection across protein-coding genes throughout genomes. Mathematically, the previously published “selection-mutation-drift model” could be theoretically used for identifying such signatures (Bulmer, 1991). However, applying this specific model to diverse species in order to identify signatures of natural selection on codon usage bias requires *a priori* empirical information about population-level genetic variations, temporal and spatial gene expression patterns, and related estimates of fitness (Yang and Nielsen, 2008; Rodrigue, Philippe and Lartillot, 2010; Tamuri, dos Reis and Goldstein, 2012; Gilchrist et al., 2015; Kubatko et al. 2016). Since such data are not available for most species, we developed an alternative statistical approach that is solely based on the analyses of DNA sequences of protein-coding genes from publicly available genomic data. In contrast to the previously published “selection-mutation-drift” model, which evaluates the relative contribution of natural selection to the observed codon usage bias by estimating selection coefficients for each possible nucleotide substitution, our approach is based on the statistical rejection of the null hypothesis that “Synonymous mutations are neutral”. Although this approach does not provide specific quantitative estimates for the level of selection on each individual gene, it serves as a robust method for identifying specific genes whose biased codon usage patterns have been impacted by natural selection.

Our statistical model is based on several key assumptions: 1) Reference genomic data represent a “wild type” genome; 2) Observed gene-specific codon usage patterns are at equilibrium; 3) For each codon, no more than one nucleotide substitution per generation is possible; 4) Nonsynonymous mutations are mostly deleterious, and therefore, the probabilities of observed nonsynonymous substitutions are negligible relative to those of synonymous ones; 5) The contributions of individual alleles of a single gene to fitness are additive; 6) Gene-specific mutational biases are independent, which allows us to ignore the possible effects of background nucleotide composition of nearby chromosomal regions on the mutational bias of focal nucleotides (Oliver et al, 2001); 7) For each individual open reading frame, the mutation rate for each of the 12 possible nucleotide substitutions (A-to-T, T-to-A, *etc.*) is constant (Rodriguez et al., 1990), which also indicates that the possible effects of adjacent nucleotides on mutation rates of each focal nucleotide could be omitted (Bérard and Guéguen, 2012); 8) The lowest mutation rate is at least 1/100 of the highest one (Drake et al., 1998).

Next, we used the previously published “selection-mutation-drift model” (Bulmer, 1991; Yang and Nielsen, 2008) to describe the relationships between the various evolutionary molecular processes that may have shaped observed gene-specific codon usage biases (Equation (1)).

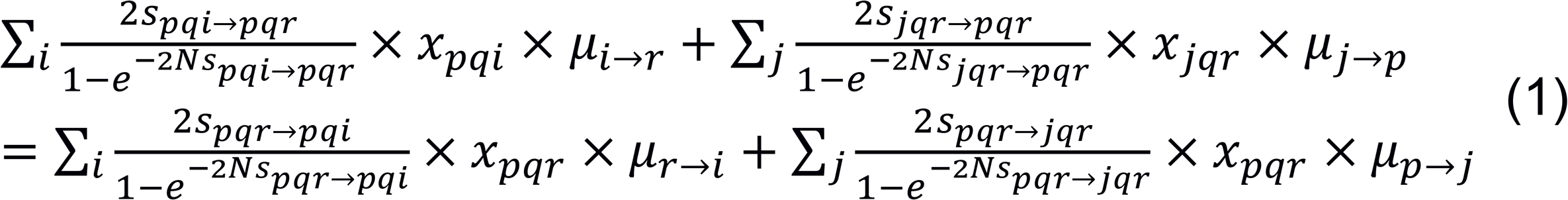

Variables *p, q*, and *r* denote the specific nucleotide identities (A, C, G, or T) of the first, second, and third positions respectively in each codon. Codons synonymous to codon *pqr* that vary at either the first or third positions are denoted by codons *jqr* or *pqi* respectively. Therefore, *S_pqh→pqr_*, for example, denotes the selection coefficient of the *pqi-to-pqr* mutation. *N* denotes effective chromosomal population size (Yang and Nielsen, 2008). *x_pqr_* denotes the count of codon *pqr* in a focal gene across the entire population, and *μ_i→r_* denotes an estimate for the *i*-to-*r* mutation rate. Consequently, the probability that an *i*-to-*r* mutation is fixed in a population is 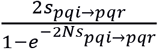 (Yang and Nielsen, 2008).

The amino acid serine represents a unique case because it is encoded by six synonymous codons that belong to two independent codon groups, which are not interchangeable via single synonymous mutations. Therefore, in our model we treat the Ser codon groups AGC, AGT and TCA, TCC, TCG, TCT as if they are coding two independent amino acids (Gilchrist et al., 2015).

Since the null hypothesis of our model specifies that all synonymous mutations are neutral, the selection coefficient of each synonymous mutation should have a zero value, which simplifies Equation (1) as follows:

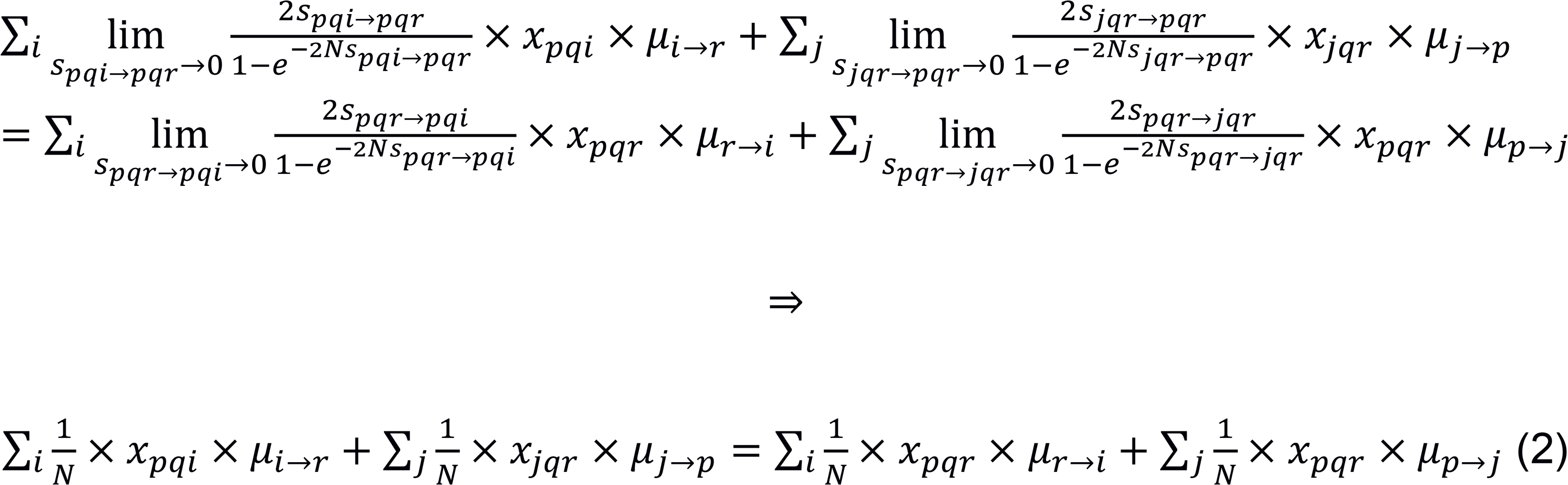

Furthermore, since each term in Equation (2) includes the factor 1, Equation (2) could be further simplified as:

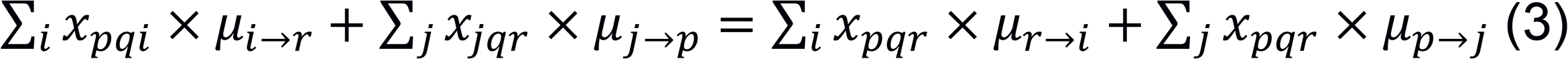

Subsequently, if the null hypothesis that all synonymous mutations are neutral is true then the impacts of genetic drift, represented by the effective chromosomal population size N, and natural selection, represented by selection coefficient s, are canceled as shown above, and therefore, the observed codon usage pattern could be explained solely by mutational bias. In contrast, if the null hypothesis is rejected then we must accept the alternative hypothesis, which indicates that natural selection has had a significant impact on the observed gene-specific codon usage bias.

To apply our derived statistical model to actual genomic data, we initially generated a dataset of expected values for gene-specific codon usage patterns across all protein-coding genes in each analyzed genome, assuming they are driven solely by mutational bias. To achieve this, Equation (3) was first used to estimate gene-specific *μ* values by using available reference genomic sequences. We then calculated expected codon counts for each gene based on the *μ* values estimated by the model. We then tested whether the model-generated (expected) and the actual (observed) gene-specific codon usage patterns are similar by using a *χ*^2^ test (df = 40) (Davis and Olsen, 2009; Xiang et al., 2015; Bera et al., 2017). Subsequently, all genes with observed codon usage patterns that were significantly different from the expected patterns generated by the model were classified as genes that carry signatures of natural selection on synonymous codon usage. As this approach only requires genetic code and the nucleotide sequences of open reading frames as input, in principle, it can be applied to any native protein-coding genes in any species whose genetic code is known.

Next, we used our approach to analyze the well-annotated genomes of five representative eukaryotic species, *Drosophila melanogaster, Arabidopsis thaliana, Saccharomyces cerevisiae, Caenorhabditis elegans*, and *Homo sapiens.* Initial analyses of the overall relative synonymous codon usage (RSCU) (Sharp et al., 1988) revealed that these species utilize markedly different genome-level patterns of synonymous codon usage biases (Fig. 1A). Next, we searched for protein-coding genes that exhibit signatures of selection on gene-specific codon usage bias by utilizing our new statistical approach. We found that after correcting for false discovery rate (FDR = 0.05)

**Figure 1.**
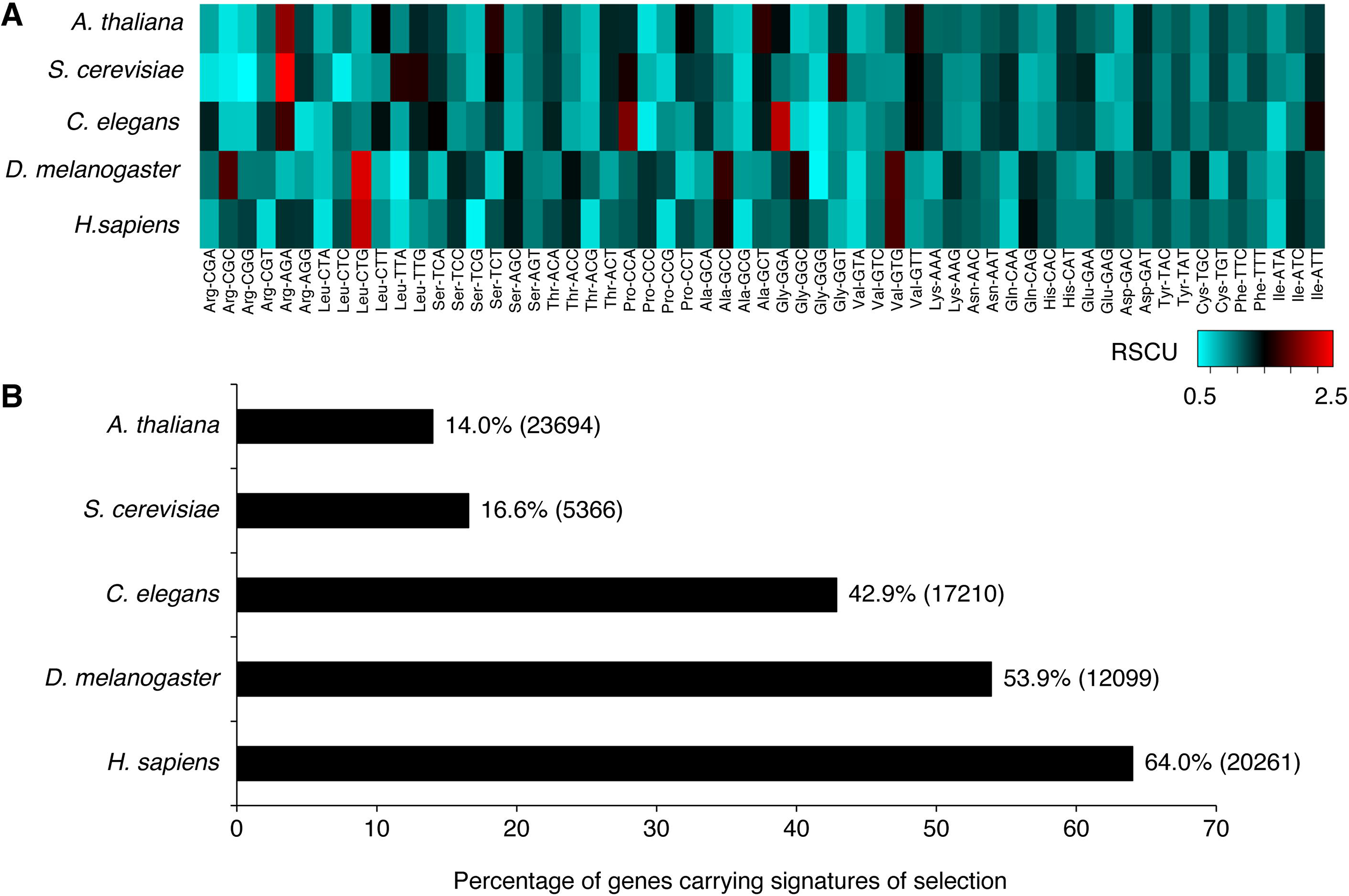
Signatures of natural selection on gene-specific synonymous codon usage bias in representative eukaryotic species. (A) Analyses of the relative synonymous codon usage (RSCU) as a measure of the overall codon usage bias across five representative Eukaryotic species. (B) Total number of protein-coding genes analyzed for each species were. *D. melanogaster*, 12099; *A. thaliana*, 23694; S. *cerevisiae*, 5366; *C. elegans*, 17210; *H. sapiens*, 20261. The estimated percentages of genes carrying selection signatures are corrected by a 0.05 false discovery rate. See also Fig. S1.

(Benjamini and Hochberg, 1995), the percentage of protein-coding genes that carry signatures of selection on gene-specific codon usage bias varied between 14% and 64% across these five representative genomes (Fig. 1B). Additional analyses of available eukaryotic genomes revealed that in 35 out of 40 eukaryotic species analyzed, at least 10% of the protein-coding genes carried signatures of natural selection on synonymous codon usage patterns, independent of whether they are unicellular or multicellular (Fig. S1). These results indicate that the overall impact of natural selection on gene-specific patterns of codon usage is broader than would be expected if synonymous substitutions were mostly functionally neutral. Our results also suggest that the relative impact of selection on gene-specific codon usage bias is not constant across the eukaryotic phylogeny, and therefore, might be the result of diverse, clade-specific selective forces.

As currently framed, our quantitative approach is based on a set of assumptions that enable the broad application of our statistical model to as many genomes as possible. However, we acknowledge that some of these assumptions might not represent the correct biological context for all species under all possible ecological conditions. We also anticipate that the assumption that adjacent nucleotides have no effect on the mutation rates of focal nucleotides could increase false-positive rates (Bérard and Guéguen, 2012). However, we anticipate that this possible source for false-positives is compensated for by the assumption that background nucleotide composition of nearby chromosomal regions (Oliver et al, 2001) is negligible, which is further alleviated by the conservative use of relaxed constraints for estimating *μ* values (Drake et al., 1998). Nevertheless, false positive rates produced by our model could be further reduced by having more precise estimates of the *μ* parameters, which could be achieved by using population-level genetic variation data, and more specific information about species-specific mutation rates, and how they might be impacted by the identities of adjacent nucleotides.

### Selection on gene-specific codon usage bias in *D. melanogaster* is associated with specific biological functions

Our statistical approach for identifying genes that carry signatures of natural selection on synonymous codon usage cannot directly address which specific synonymous codons might be under selection in each selected gene. Therefore, we next used a clustering approach to determine whether the genes we have identified in our initial screen share similar codon usage patterns, or alternatively, represent a heterogeneous population comprised of multiple gene clusters, each defined by a unique pattern of biased codon usage.

Due to its well-annotated genome and wealth of available genetic and phenotypic data, we chose to use the genetically tractable species *D. melanogaster* for functional analyses of genes that are under selection for biased codon usage patterns. Hierarchical gene-clustering analysis revealed that genes with identified signatures of selection on biased codon usage could be assigned to one of two major clusters (Clusters A and B; Fig. 2). Further analysis of Cluster A revealed that synonymous codon usage in these genes is enriched for the rare codons Lys-AAA, Glu-GAA, Gln-CAA, Phe-TTT, Tyr-TAT, and His-CAT, relative to their usage in Cluster B genes. Similar clustering analyses of the genomes of *A. thaliana, S. cerevisiae, C. elegans*, and *H. sapiens* revealed gene-by-codon clustering in all five species, likely representing species-specific patterns of selection (Figs. S2-5).

**Figure 2.**
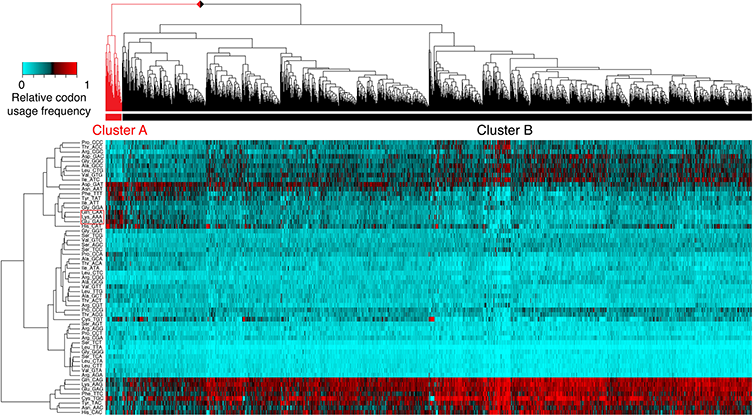
Clustered codon usage patterns across genes carrying signatures of selection on gene-specific codon usage bias in *D. melanogaster.* Hierarchical clustering of all genes (horizontal axis) identified as carrying significant signatures of selection on gene-specific codon usage bias. The relative usage frequencies of each codon in each gene are color-coded (vertical axis). Euclidean distances were used to measure dissimilarities between codons. Spearman’s correlation coefficients were used to measure similarities between genes. Complete linkage was used as the clustering criterion. Clusters A (red bar) and B (black bar) represent the two major gene groups identified. See also Figs. S2-5, Table S1.

Next, we tested the hypothesis that the identified gene clusters in *D. melanogaster* represent specific biological functions. Analyses of previously published tissue-specific gene expression data (Chintapalli, Wang, and Dow, 2007) revealed that genes that are preferentially expressed in the male accessory glands are significantly overrepresented in Cluster A and underrepresented in Cluster B (Table 1). Similarly, analysis of previously published data on sexually dimorphic genes (Graveley et al., 2010; Gelbart and Emmert, 2013) revealed that genes with male-enriched expression are significantly overrepresented in Cluster A and underrepresented in Cluster B (Table 2). These data suggest that a specific combination of rare codons contributes to the function of some genes associated with the male reproductive system.

**Table 1.** Tissue-specific expression patterns of genes in Clusters A and B. Genes included in Clusters A and B are as in Fig. 2. Significant over-or under-representation is shown in bold. N/A means that no tissue-specific genes were found in the entire genome or the focal gene cluster. Although present in the FlyAtlas database, we did not identify tissue-enriched genes in the thoracic-abdominal ganglion, virgin female spermatheca, inseminated female spermatheca, and adult fat body, and therefore, are not included here. Fold enrichment is relative to the entire genome. Bonferroni correction is applied, p< 0.05/20 = 0.0025. See also Table S1.

|  | Cluster A |  | Cluster B |  |
| --- | --- | --- | --- | --- |
| Tissue | Fold enrichment | p-value | Fold enrichment | p-value |
| Adult Central Nervous System | 0.48 | 0.378 | 1.13 | 0.04 |
| Brain | N/A | N/A | 0.57 | 0.205 |
| Crop | N/A | N/A | 1.15 | 0.622 |
| Midgut | 2.31 | 0.066 | 1.19 | 0.003 |
| Hindgut | N/A | N/A | 1.01 | 0.582 |
| Malpighian Tubules | 1.11 | 0.599 | 1.1 | 0.205 |
| Ovary | 1.71 | 0.256 | 1.09 | 0.123 |
| Testis | 1.36 | 0.082 | **0.56** | **<0.001** |
| Male Accessory Gland | **4.45** | **<0.001** | **0.36** | **<0.001** |
| Carcass | N/A | N/A | **0.34** | **<0.001** |
| Salivary Gland | N/A | N/A | 0.49 | 0.024 |
| Heart | N/A | N/A | 0.86 | 0.493 |
| Eye | 2.53 | 0.187 | 0.8 | 0.05 |

**Table 2.** Sex-biased expression patterns of genes in Clusters A and B. Genes included are as in Table 1. Significant over- or under-representation is shown in bold. Fold enrichment is relative to the entire genome. Bonferroni correction is applied, p< 0.05/4 = 0.0125. See also Table S1.

| Gene cluster | Sex | Fold enrichment | p-value |
| --- | --- | --- | --- |
| Cluster A | Male | **1.66** | **<0.001** |
|  | Female | 1.67 | 0.091 |
| Cluster B | Male | **0.61** | **<0.001** |
|  | Female | 1.07 | 0.060 |

In addition, analyses of Gene Ontology terms (Gene Ontology Consortium, 2015) revealed that genes that encode extracellular matrix proteins are overrepresented, and genes that encode cytoplasmic proteins are underrepresented, in Cluster A (Table S1). We also found that genes annotated as encoding odorant-binding proteins, a class of secreted proteins, as well as extracellular space proteins, are underrepresented in Cluster B. Together, these analyses suggest that in *D.* melanogaster, natural selection on the increased usage of specific combinations of rare synonymous codons might be important for male-specific functions of some genes.

### Biased gene-specific codon usage can modulate spatial protein expression patterns

One possible explanation for the association between certain biased gene-specific codon usage patterns and tissue-specific expression is that the selected specific rare synonymous codons match tissue-specific tRNA pools, and therefore, enhances the translation efficiency of some protein in specific tissues or cell types (Zalucki, Gittins and Jennings, 2008; Pechmann, Chartron and Frydman, 2014). Similar associations between gene-specific codon usage bias and organ-enriched gene expression patterns have been previously reported in humans (Plotkin, Robins, and Levine, 2004; Dittmar, Goodenbour and Pan, 2006). Therefore, we next tested the hypothesis that the specific pattern of codon usage bias exhibited by Cluster A genes contributes to increased translation efficiency in the male reproductive system.

Our clustering analysis revealed that male-enriched genes in Cluster A preferentially use the rare codons Lys-AAA, Gln-CAA, Glu-GAA, Phe-TTT, Tyr-TAT, and His-CAT (Fig. 2). However, only the first three codons are recognized by exactly matched tRNA anticodons, while the other three rare codons share the same tRNAs with their more commonly used synonymous codons, with a mismatch (wobble) at the third codon position (Crick, 1966; Chan and Lowe, 2009), which suggests that these codons are less likely to contribute to tissue-specific translation efficiency. Therefore, we next analyzed the specific usage of rare codons AAA, CAA, and GAA in all genes that show enriched expression in the male reproductive system, independent of whether these genes have passed the initial statistical threshold for the detection of selection on gene-specific codon usage patterns. We found that these specific rare synonymous codons are indeed overrepresented in male-reproductive-system-specific genes (Table 3). Thus, not only are genes preferring AAA, CAA, and GAA more likely to be male-reproductive-system-specific, but also genes specifically transcribed in male reproductive system use AAA, CAA, and GAA more often than their expected genome average usage.

**Table 3.**
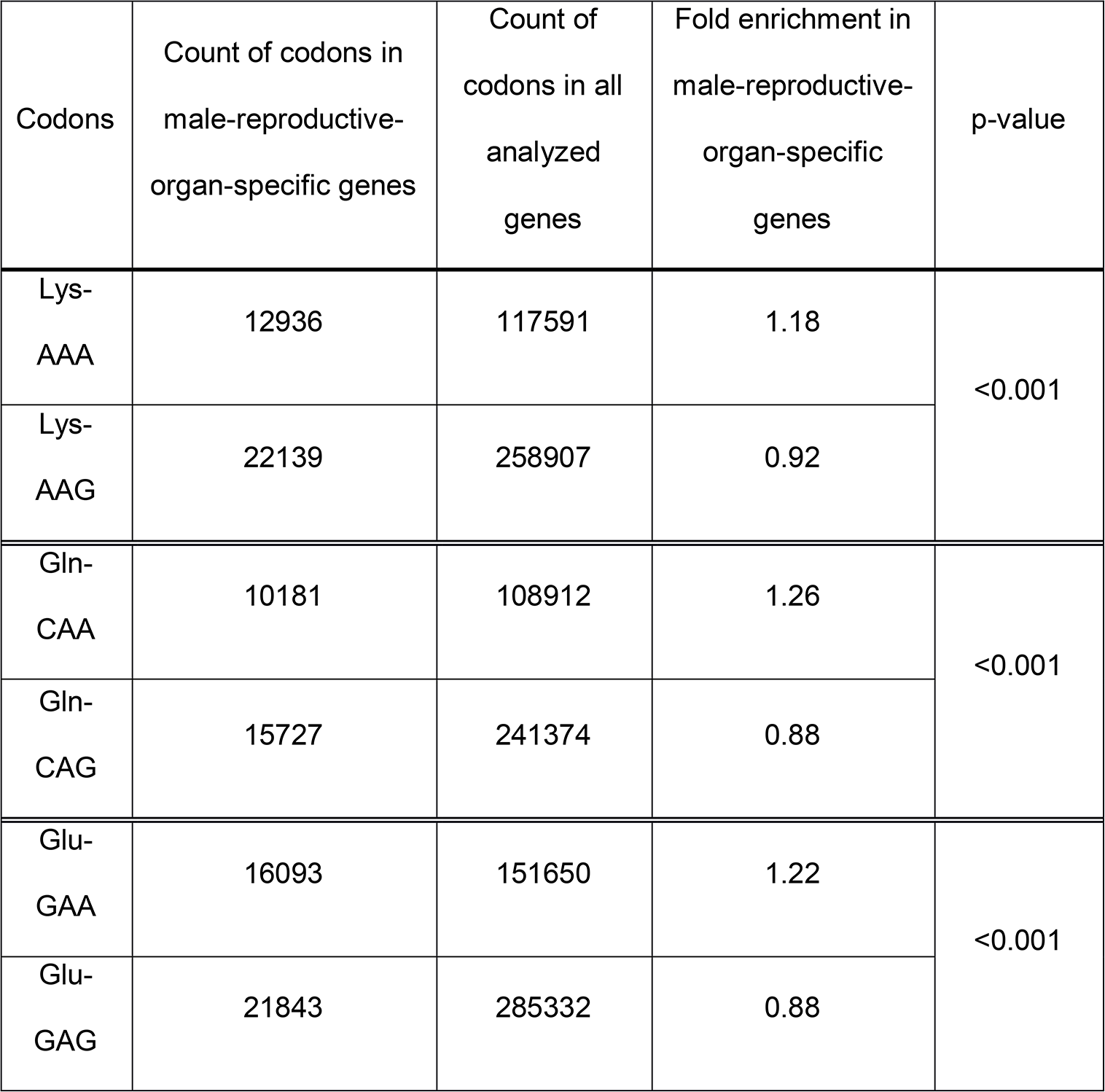
Usage frequencies of rare codons Lys-AAA, Gln-CAA, and Glu-GAA in the male reproductive tissues relative to whole genome averages. Counts of relevant codons were calculated for the entire genome and male-reproductive-organ-specific genes (including genes with enriched expression in the testis and the accessory-glands). Hypergeometric tests were used to test whether a specific synonymous codon is over-or under-represented in male-reproductive-organ-specific genes. Fold enrichment is calculated by dividing the relative frequency of a focal codon in male-reproductive-organ-specific genes by the relative frequency of the focal codon usage across all protein-coding genes.

If the selection on an increased relative usage of rare codons AAA, CAA, and GAA serves a biological function that is specific to gene function in the male reproductive system then it should depend, at least in part, on the increased expression of the specific rare tRNAs that bind these specific codons. In accordance with this hypothesis, northern blot analyses of tissue-specific tRNA expression revealed that tRNA^Lys^_TTT_ is enriched in the male reproductive organs, further supporting the hypothesis that spatial regulation of some tRNA genes that correspond to rare codons could contribute to tissue-specific increase in protein translation efficiency (Fig. 3A-C). The expression patterns of the tRNAs that match Glu-GAA and Gln-CAA were not investigated because their sequences are almost identical to their common tRNA counterparts, which does not allow their independent detection by hybridization probes (de Muro, 2008). Nevertheless, these data indicate that some *Drosophila* genes have evolved a specific codon usage bias that specifically drives their levels and/or activity in the male reproductive system.

**Figure 3.**
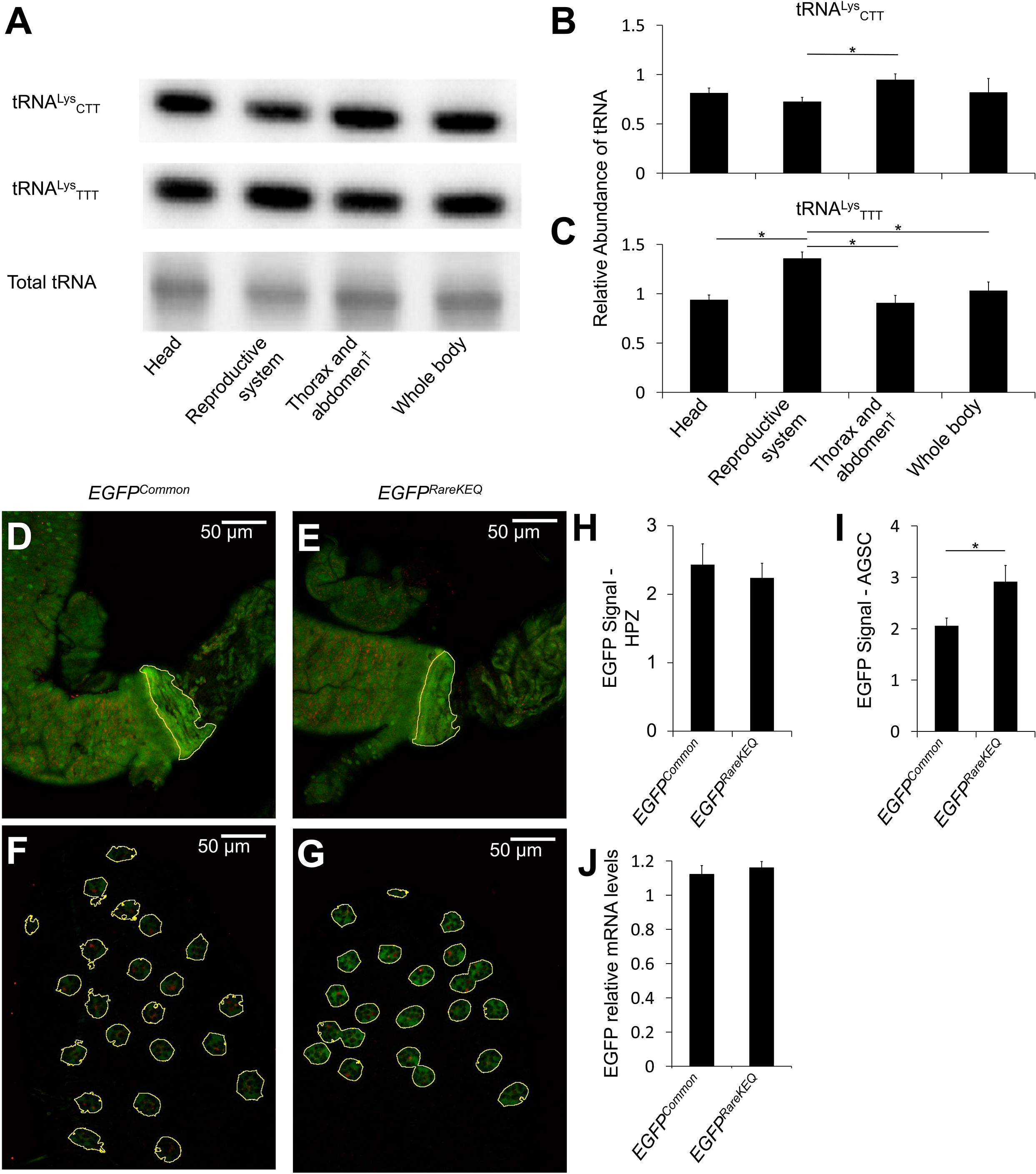
Gene-specific codon usage bias affects spatial protein expression in *D. melanogaster.* (A) Representative northern blots and (B-C) summary data of the relative abundance of tRNA^Lys^_CTT_ (common) and tRNA^Lys^_TTT_ (rare) codons across male tissues. *, p<0.05 (n = 4, ANOVA followed by SNK *post hoc* tests). Error bars denote standard deviation. |: Reproductive system is excluded. (D-E) Representative images showing EGFP and RFP expression in the HPZ (yellow lines mark tissue boundaries). (F-G) Representative images showing EGFP and RFP expression in the AGSC (yellow lines mark secondary cells). (H-I) Summary data of normalized EGFP signals in HPZ and AGSC. *, p<0.01 (n=5, two-tailed unpaired Student’s t-test). Error bars denote standard deviation. (J) Real-time qRT-PCR mRNA expression data of *EGFP^Common^* and *EGFP^RareKEQ^* (two-tailed unpaired Student’s t-test, n = 4, NS). Error bars denote SEM.

Next, we empirically tested whether specific combinations of rare codons are sufficient to drive tissue-specific translation efficiency by generating transgenic flies that express different alleles of the EGFP reporter under the control of the UAS-GAL4 binary expression system (Brand and Perrimon, 1993). One allele was comprised of common codons for all amino acids (*EGFP^Common^*), and the second allele was comprised of rare codons AAA, GAA, and CAA for amino acids Lys, Glu, and Gln respectively (*EGFP^RareKEQ^*), which represent about 18% of the total residues in EGFP. Each *EGFP* allele was also co-expressed with an *RFP* reporter encoded by common codons, which served as a transgene expression control. Both transgenes were driven by the ubiquitous Act5C-GAL4 driver (Kopp, Blackman, and Duncan, 1999). To evaluate the effect of codon usage bias on the spatial pattern of both EGFP alleles *in vivo*, we compared the two genotypes by measuring the RFP-normalized EGFP signals in the accessory gland secondary cells (AGSC) and in the hindgut proliferation zone (HPZ) (Takashima et al., 2013). We found that the signal from the *EGFP^RareKEQ^* allele in the AGSCs was significantly higher than that of the *EGFP^Common^* allele. In contrast, both alleles produced similar signals in the HPZ (Fig. 3D-I). Because the overall transcript levels of both alleles are similar (Fig. 3J), these results suggest that some specific combinations of synonymous rare codons could enhance protein translation in the *Drosophila* male reproductive system.

## DISCUSSION

In contrast to a broadly accepted dogma in evolutionary biology and population genetics, our theoretical and empirical studies indicate that synonymous mutations should not be considered neutral in general. By using multiple complementary approaches, we demonstrate here that the impact of natural selection on gene-specific codon usage bias is common across the eukaryotic phylogeny and is likely to play a much more prominent role in defining gene-specific codon usage bias than was previously assumed. Furthermore, our data indicate that in multicellular organisms such as the fruit fly, natural selection is likely to optimize codon usage patterns to match specific spatial and temporal constraints on gene function, and therefore, what might represent an optimal codon usage for individual genes does not necessarily mean common codons should be expected at every position. Our findings also indicate that previous reports about the phenotypic consequences of synonymous mutations are not anecdotal rare cases. Instead, these cases likely represent a fundamental biological principle associated with the spatial and temporal gene activity landscapes in eukaryotes. In addition, our analyses of gene-specific codon usage biases in *D. melanogaster* suggest that gene-specific codon usage bias is heterogeneous across protein-coding genes in a single species, and therefore, likely to be the outcome of diverse selective pressures associated with different biological functions. Furthermore, our data suggest that different classes of rare synonymous codons exhibit differential sensitivities to natural selection.

Consequently, our discovery that gene-specific codon usage patterns are often shaped by natural selection indicates that alternative synonymous codons serve important, yet often overlooked, biological functions. Although several previous studies have used statistical approaches to suggest that synonymous mutations in specific genes and species might not be neutral (Berger, 1977; Eyre-Walker, 1991; Stenico, Lloyd, and Sharp, 1994; Lawrie et al., 2013), our current study provides the first genome-scale comprehensive view of the overall impact of natural selection on gene-specific synonymous codon usage patterns across diverse eukaryotic taxa. Thus, our unbiased and broadly applicable statistical approach has uncovered a fundamental genomic organizational principle that is sensitive to natural selection, and likely represents an important component of the genotype-phenotype axis in diverse eukaryotic species and biological contexts. Therefore, it is likely that investigations, which rely on the assumption that synonymous mutations are neutral, often overestimate or underestimate the prevalence of selection on specific protein-coding genes, which could impact the interpretations of the role of these genes in genotype-phenotype relationships at the level of populations and the evolutionary timescale in particular (Chen et al., 2010), and the genome evolution in general (Campos et al., 2012; Matsumoto et al., 2016). Specifically, the prevailing assumption that most synonymous mutations are neutral, and therefore represent a robust and quantitative measure of selection-independent evolutionary rates of nucleotide substitutions, needs to be revised. This is especially important in the context of the commonly used tests for identifying molecular signatures of selection in specific protein-coding genes, and for classifying the modes of natural selection (*e.g.*, “positive” versus “negative” selection) (McDonald and Kreitman, 1991; Nielsen, 2001; Echave, Spielman, and Wilke, 2016).

Our data also indicate that, similarly to *Drosophila*, many human protein-coding genes carry signatures of selection on their biased codon usage patterns (Fig. 1). This finding could have broad implications for studies of genetic variants underlying quantitative phenotypes in human populations. Consequently, assuming that synonymous mutations are functionally neutral could lead to a biased view that associations between nonsynonymous SNPs and specific phenotypic values are the only ones important for understanding overall trait variations (Hampe et al., 2007; WTCCC and TASC, 2007; Xu and Taylor, 2009; Fong et al., 2010; Huang et al., 2012). This is supported by a recent data survey of human GWAS data from over 2000 independent studies, which suggested that synonymous and nonsynonymous SNPs have similar likelihoods and effect sizes in terms of association with disease phenotypes (Chen et al., 2010).

Although our analysis of human gene-specific codon usage bias was based on data from a single reference genome, the simplest interpretation of our results is that both synonymous and nonsynonymous mutations are likely influenced by natural selection in human populations, and therefore, many synonymous SNPs could still have significant quantitative and/or qualitative impacts on gene functions and associated phenotypes in health and disease.

We also found that gene-specific codon usage bias is not uniform across the genome. Instead, our data suggest that different groups of genes in the same species often exhibit enrichment for specific combinations of rare or common codons, which suggests that different combinations of synonymous codons may have been selected for independent biological reasons. In this regard, we show that at least one group of genes with enriched expression in the reproductive system of male *Drosophila* preferentially use the rare codons Lys-AAA, Gln-CAA, and Glu-GAA. By using allelic variants of a reporter gene, we show that the selective usage of these three rare codons is sufficient for generating spatially biased protein expression patterns. These findings are consistent with some of the assumptions about the possible roles of codon usage bias, first put forward by the “translational selection theory” (Ikemura, 1981; dos Reis, Savva, and Wernisch, 2004; Plotkin, Robins, and Levine, 2004; Dittmar, Goodenbour, and Pan, 2006). Consequently, our study provides a broader context to this fundamental evolutionary theory by emphasizing the possible role of gene-specific codon usage bias in the spatial regulation of proteins in multicellular eukaryotes. However, regulation of spatial protein expression cannot explain all cases of selection on gene-specific codon usage bias because genomes of some unicellular eukaryotes, such as the choanoflagellate *Monosiga brevicollis* and the green algae *Chlamydomonas reinhardtii*, also include a significant proportion of protein-coding genes that exhibit signatures of selection on their codon usage patterns (Fig. S1).

In addition to providing a fundamental aspect of gene regulation, our discovery that the impact of natural selection on gene-specific codon usage bias is prevalent in eukaryotic genomes also carries important practical implications for experimental systems that depend on heterologous gene expression. Specifically, we have found that for some genes, the selective usage of synonymous rare codons could increase their translation efficiency in specific tissues and cell types. Therefore, designing experiments with the assumption that codon optimization of a transgene necessitates the preferential usage of common synonymous codons could lead to unexpected phenotypic outcomes, and could affect data interpretation, especially for genes with restricted spatial and/or temporal expression patterns.

Our studies also highlight the need for a better understanding of the mechanisms that regulate the spatial expression patterns of tRNAs. Some previous studies have argued that tRNA gene copy number is likely the primary mechanism that regulates the relative levels of individual tRNAs in cellular pools (dos Reis, Savva and Wernisch, 2004). Yet, variations in gene copy number alone cannot explain the tissue-specific expression patterns of some rare tRNAs observed by us (Fig. 3A-C) and others (Dittmar, Goodenbour and Pan, 2006; Gingold et al., 2014). Therefore, although tRNAs are thought to be exclusively transcribed by the constitutively-active RNA Pol III complex, there must be additional molecular mechanisms that enable the temporal and spatial regulation of some tRNAs at the transcriptional and/or post-transcriptional levels (Yoshihisa et al., 2007; Wichtowska, Turowski, and Boguta, 2013; Turowski and Tollervey, 2016).

Although our empirical work here has mostly focused on a specific set of rare codons that bind unique tRNA anti-codons, it should be noted that genes in Cluster A (Fig. 2) also preferentially use the rare codons Phe-TTT, Tyr-TAT, and His-CAT. These rare synonymous codons share their tRNA anticodons with their more common synonymous codons via a wobble mechanism (Chan and Lowe, 2009). Theoretical models and empirical data suggest that this specific class of rare codons is likely to slow down translation due to the lower affinity of the common tRNAs to their associated rare synonymous codons (Ikemura, 1981; dos Reis, Savva, and Wernisch, 2004; Savir and Tlusty, 2013; Brule and Grayhack, 2017). Therefore, we speculate that in contrast to rare codons that bind unique perfectly-matching tRNAs, rare codons that share tRNAs with their more common synonymous codons may have evolved to broadly restrict translation efficiency of some low-abundance proteins, or proteins that require slow folding kinetics associated with specific post-translation modifications, or the binding of other protein and small-molecule partners, chaperones, or co-factors (Komar, Lesnik and Reiss, 1999; Weygand-Durasevic and Ibba, 2010; Angov, 2011; Zhou et al., 2013; Fluman et al., 2014; Pechmann, Chartron and Frydman, 2014; Chaney and Clark, 2015; Quax et al., 2015; Yu et al., 2015; Fu et al., 2016; Zhao, Yu, and Liu, 2017).

In summary, the results we present here suggest that in contrast to a widely accepted dogma, synonymous mutations should not be considered functionally and evolutionarily neutral by default. Instead, as was previously shown for a few specific protein-coding genes (Hasegawa, Yasunaga, and Miyata, 1979; Ikemura, 1981; Akashi, 1994; Stenico, Lloyd, and Sharp, 1994; Ko et al., 2005; Pagani, Raponi, and Baralle, 2005; Kimchi-Sarfaty et al., 2007; Parmley and Hurst, 2007; Zhang, Hubalewska, and Ignatova, 2009; Gu et al., 2012; Zhou et al., 2013; Ma et al., 2014; Shin et al., 2015; Fu et al., 2016), our data indicate that selection on synonymous codon usage is likely to serve a more general function in diverse biological processes across Eukaryotes. Nevertheless, although our data indicate that “synonymous” and “neutral” mutations often represent different biological values in the context of the genotype-phenotype axis, we do not suggest that the concept of evolutionary neutrality, first described by Darwin in the *On the Origin of Species* as *“Variations neither useful nor injurious would not be affected by natural selection”*, should be overall rejected. Instead, we argue that the commonly accepted specific assumption that all synonymous mutations are generally neutral should be reevaluated.

## ACKNOWLEDGEMENTS

Z.P. dedicates this manuscript to Dr. Yang Zhong of Fudan University, a prominent botanist and evolutionary biologist who had a great influence on his choice to pursue a research career in mathematical and evolutionary biology. The authors also wish to thank Alan Templeton, Allan Larson, and members of the Ben-Shahar and Zaher laboratories for helpful discussions. They also thank Paula Kiefel, Dianne Duncan, and Ian Duncan for technical assistance with generating transgenic animals and microscopic imaging. Z.P. is supported by the NIH training grant 2T32HG000045. This research has been supported in part by NIH grant NS089834, and NSF grants 1545778 and 1707221 to YB, and NIH grant 5R01GM112641 to HZ.

## AUTHOR CONTRIBUTIONS

ZP and YB conceived the study; ZP performed the studies under the guidance of YB and HZ; HZ assisted with the design and execution of molecular biology studies; ZP, HZ, and YB wrote the manuscript and approved its final version.

## EXPERIMENTAL PROCEDURES

### Genomic and transcriptomic data

Protein-coding DNA sequences were from Ensembl (Zerbino et al., 2017) (https://www.ensembl.org/). Reference coding sequences included in the analysis of gene-specific codon usage bias were chosen according to the following criteria: 1) The sequence length is a multiple of three. 2) The sequence uses standard genetic code. 3) For each gene, only the longest mRNA isoform was used for analysis. If there were multiple isoforms of the same length, then the first record shown in the fasta file was used. 4) The encoded protein includes all 19 amino acids that have degenerate codons; for the amino acid serine, the two-fold and four-fold degenerate codon groups were treated as if they encoded two different amino acids. Transcriptomic data were from the FlyAtlas microarray database (Chintapalli, Wang, and Dow, 2007) and the modENCODE RNA-seq database (Graveley et al., 2010; Gelbart and Emmert, 2013).

### Estimating *μ* and expected codon counts

The relationships between *μ* values and codon counts are described by Equation (3) (Results section). Based on the standard genetic code and all possible combinations for synonymous nucleotide substitutions, we classified all degenerate codons into six categories. For each category, we used Equation (3) to generate a homogeneous linear equation system. Subsequently, we treated one of the x variables as a known parameter and analytically solved all other x variables. Finally, these solutions were used to calculate the expected codon counts of a protein-coding gene.

Category one – Codons for Arginine.

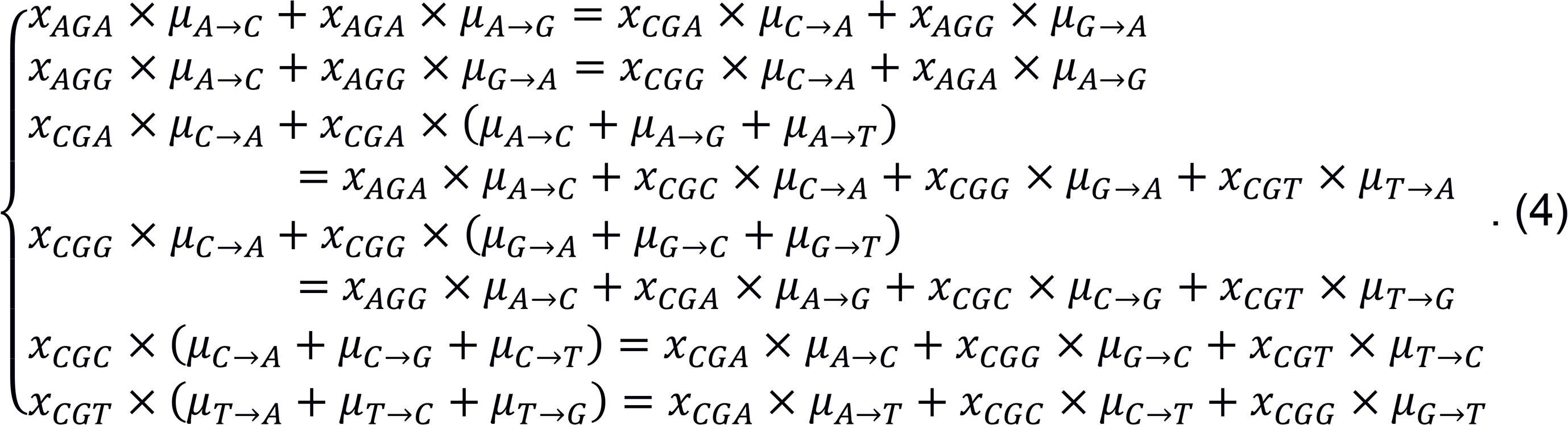

Category two – Codons for Leucine:

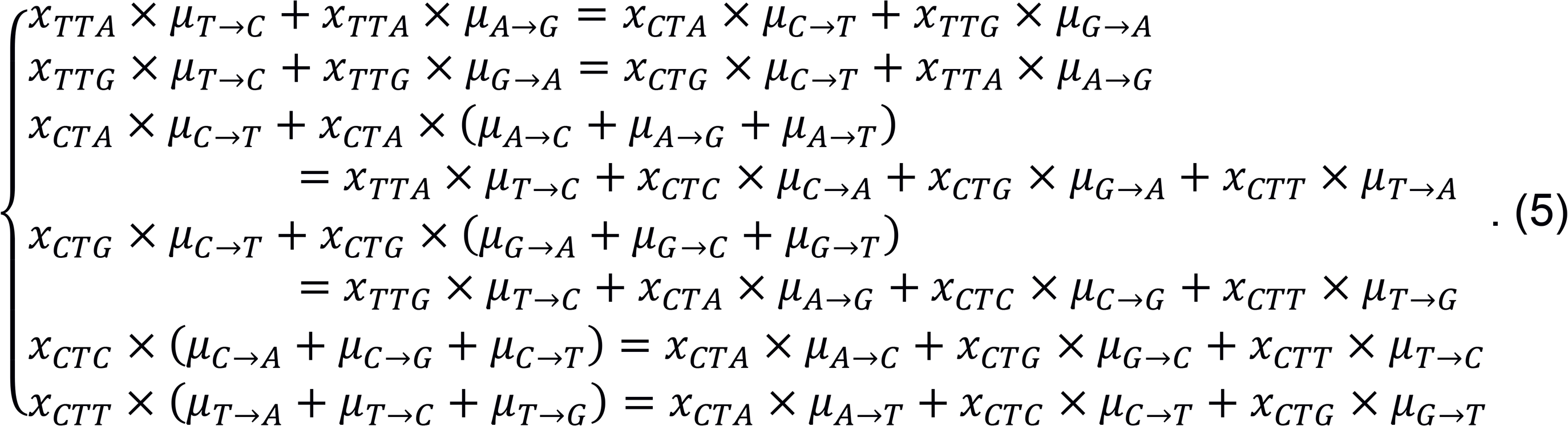

Category three – Four-fold degenerate codons:

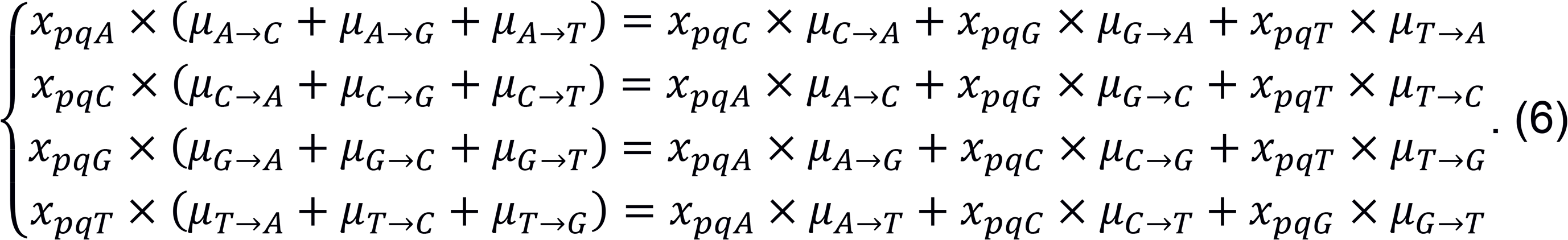

Category four – Codons for Isoleucine:

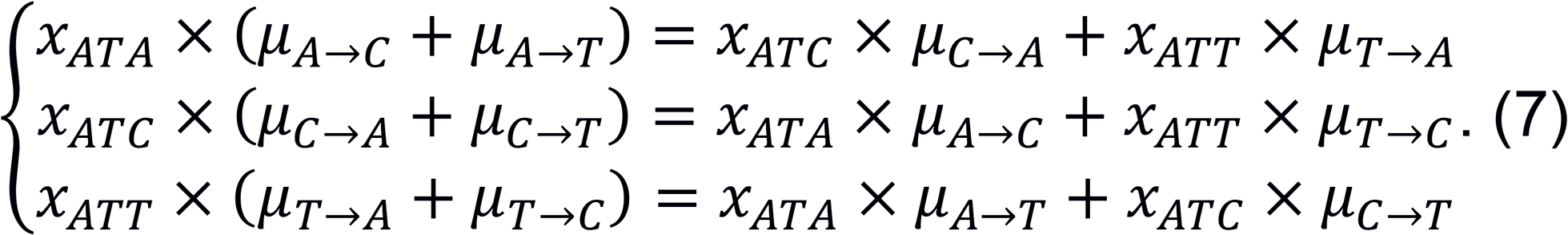

Category five – C/T-ended two-fold degenerate codons:

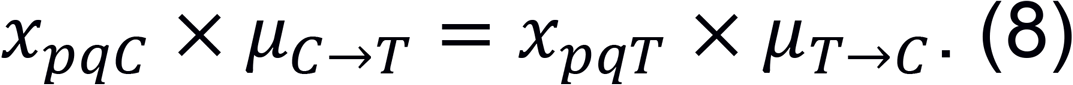

Category six- A/G-ended two-fold degenerate codons:

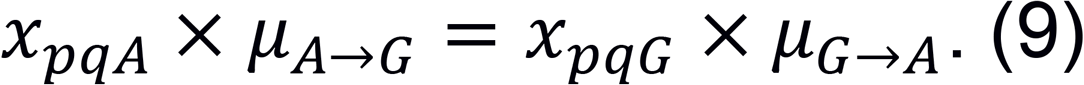

Information about the analytical solutions of the above equations is shown in the Supplemental Information (Analytical Solutions of Equation Systems Used to Estimate *μ* Values). Using these solutions, with a given set of *μ* values and counts of amino acid residues, we can calculate the expected counts of synonymous codons. For example, the expected counts of codons for Lys can be calculated by

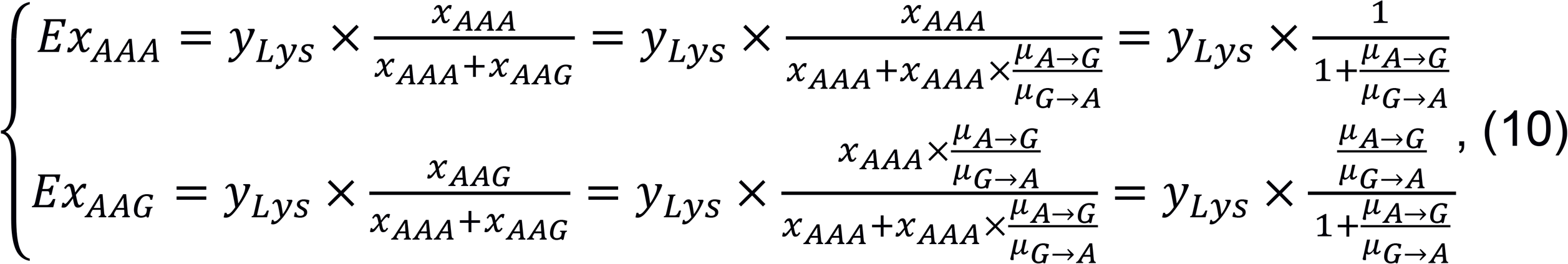

where *EX_AAA_* is the expected count of AAA codon, and *y_Lys_* is the count of Lys residues.

Then we can use the expected counts and the observed real counts of all degenerate codons to calculate a *χ*^2^ value. Since for a given protein-coding sequence, the *χ*^2^ value is a function of *μ* values, we can define this function as *χ*^2^(Θ), where *Θ* is a vector describing all *μ* values,

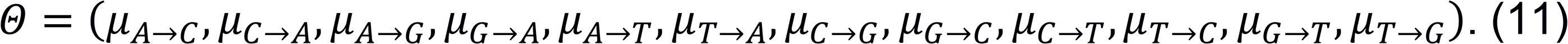

For a gene, to estimate *μ* values, we try to minimize the value of *χ*^2^(Θ) by changing the elements in Θ, using the sequential least squares programming (SLSQP) algorithm (Kraft, 1988).

Since equations generated from Equation (3) form a system of homogeneous linear equations, it is meaningless to infer exact *μ* values from the minimization of *χ*^2^(Θ); rather, only the relative magnitudes of different *μ* values are important for calculating the expected codon counts later. As we used the assumption that the lowest *μ* value is at least 1/100 of the highest, we set the range of *μ* values between 0.001 and 0.1 during the minimization of *χ*^2^(Θ). For each gene, the minimum *χ*^2^(Θ) is used to calculate p-value. Since we assumed that reference genomic data represent a “wild type” genome, the procedure mentioned above was applied to each individual reference gene.

### Codon usage heatmaps

For a protein-coding gene g, the relative usage frequency *f_gd_* of a codon *d* is defined as

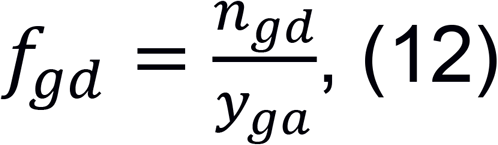

where *n_gd_* is the count of *d* in *g*, and *y_ga_* is the count of amino acid *a* encoded by *d* and its synonymous codons in g. It should be noted that codons for Ser are not treated as two codon groups in codon usage heatmaps. As the mechanism of recognizing stop codons is fairly different from recognizing other codons (Brown et al., 2015), and methionine and tryptophan respectively have only one codon, the analysis is restricted to the other 59 codons. Therefore, a 59-dimension vector *B_g_* is used to describe the codon usage pattern of *g*,

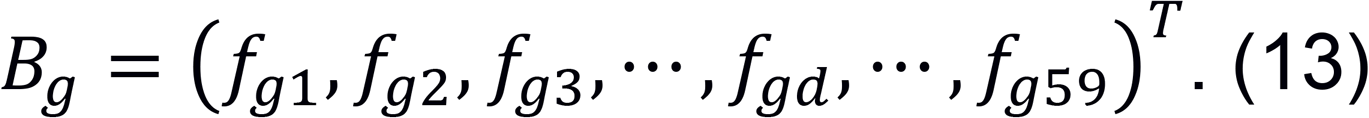

For a genome containing *M* protein-coding genes, an M-dimension vector *H_d_* is used to describe how often a codon *d* is used across these genes,

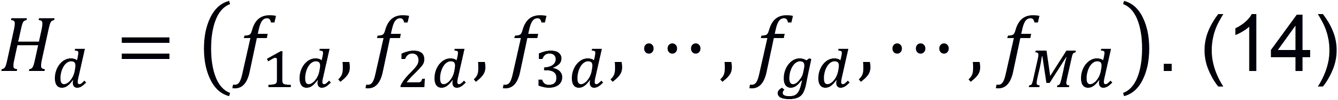

Both *B_g_’s* and H_d_s are hierarchically clustered. The values of all *f_gd_*’s are then color-coded to generate a codon usage heatmap.

The “gplots” (Warnes et al., 2015) package in R was used to generate heatmaps. The method of hierarchical clustering was complete linkage with Euclidean distance measuring the dissimilarities between codons and Spearman’s correlation coefficient measuring the similarities between genes.

### Identifying genes with tissue-specific expression patterns

Mean adult gene expression data from the FlyAtlas database were used to identify genes with tissue-specific expression patterns. A gene was classified as tissue-specific if its average mRNA level in a specific tissue was at least ten-fold to the tissue with the second highest mRNA level. To reduce redundancy, “head” expression values were excluded from the analysis. Specifically, since “brain” and “thoracic-abdominal ganglion” both belong to the central nervous system (CNS) and many CNS-specific genes are expressed in both places, “adult central nervous system”-specific mRNA level is represented by the higher mRNA level between “brain” and “thoracic-abdominal ganglion”. When “adult central nervous system” was involved in the analysis, the “brain” and “thoracic-abdominal ganglion” were excluded, and *vice versa.*

### Identifying genes with sex-biased expression patterns

Adult fly expression data from the modENCODE RNA-seq database were used to identify genes with either male- or female-biased expression patterns. Only genes that showed at least ten-fold expression in one sex relative to the other were defined as sex-biased.

### Hypergeometric tests for gene set enrichment

The hypergeometric tests were performed using the online tool at http://www.rothsteinlab.com/tools/apps/hyper_geometric_calculator.

### Gene ontology analysis

Gene ontology (GO) analysis (Gene Ontology Consortium, 2015) of *Drosophila* genes was performed by using online tools (http://geneontology.org/). Category enrichments were determined by comparing term frequencies between each gene cluster and the whole genome, followed by a Bonferroni correction.

### Animals

Fruit flies (*D. melanogaster*) were raised on corn syrup-soy food (Archon Scientific) at 25°C and 60% relative humidity with a 12-hour light/dark cycle. Synthetic cDNAs encoding EGFP and mCherry fluorescent proteins were from IDT Inc. Full sequences of *mCherry^Common^*, *EGFP^Common^*, and *EGFP^RareKEQ^* alleles are in the Supplemental Information (Sequences of Fluorescent Reporter cDNAs). To generate animals that express each transgenic allele under UAS control, synthetic cDNAs were cloned into the EcoRI/NotI cloning sites of the pUASTattB plasmid (Bischof et al., 2007). Since the *EGFP^RareKEQ^* allele contains one EcoRI site in the middle, digestion time was shortened to less than 20 minutes to allow incomplete digestion. The *UAS-RFP^Common^* transgene was inserted into a chromosome II landing site (Bloomington #24483), and the UAS-*EGFP^Cĳmm¤n^* and *UAS-EGFP^RareKEQ^* transgenes were inserted into the same chromosome III landing site (Bloomington #24749). Double homozygotic lines *UAS-RFP^Common^;* UAS-*EGFP^Common^* and *U AS-RFP^Common^; UAS-EGFP^RareKEQ^* were generated and then crossed to

*Act5C-GAL4/SM6* (Duncan lab, Washington University). Canton-S isoA flies were used for tRNA northern blot.

### Analyses of tRNA gene expression analyses

Northern blots were used to measure relative tRNA abundance in different body parts. Total RNA was extracted from four pools of 10 dissected male tissues (head, reproductive system, and remaining thorax and abdominal parts) and 10 whole male flies with the TRIzol reagent (Invitrogen Catalog # 15596-026). Probe sequences were. tRNA^Lys^CTT, 5’-AACGTGGGGCTCGAACCCACGACCCTGA-3’; tRNA^Lys^TTT, 5’-GAACAGGGACTTGAACCCTGGACCCTTG-3’. Probes were labeled by ^32^P using T4 PNK (NEB Catalog # M0201S). Signals were measured and normalized to total tRNA signals using the BIO-RAD Quantity One 1-D analysis software. ANOVA followed by SNK *post hoc* test was used to compare tRNA levels between samples.

### Real-time qRT-PCR

The mRNA expression levels of reporter genes in whole four-day-old male flies were quantified by using real-time qRT-PCR, following previously published methods (Lu et al., 2012; Hill et al., 2017). For RpL32, the forward primer was 5’-CACCAAGCACTTCATCCG-3’, and the reverse primer was 5’-TCGATCCGTAACCGATGT-3’. For *EGFP^Common^* and *EGFP^RareKEQ^*, the forward primers were respectively 5’-AACTTCAAGATCCGCCACAAC-3’ and 5’-AACTTCAAAATCCGCCACAAC-3’, while these alleles shared the same reverse primer 5’-GTGCTCAGGTAGTGGTTATCG-3’.

### Quantitative reporter gene imaging

Male reproductive and gut tissues from four-day-old adult male flies that express either the *EGFP^Common^* or *EGFP^RareKEQ^* were dissected in chilled PBS and mounted for imaging on a Nikon A1Si laser scanning confocal microscope with a 20X oil objective (n=5 samples per genotype). All images were taken within 10 minutes of dissection. Single plane fluorescent images of the AGSC were used to estimate EGFP expression levels of each allele. Similar images of the HPZ were used as generic tissue controls. The NIS-Element Ar software was used to capture EGFP and RFP signals and generate channel-merged images. Normalized EGFP signals from each image were quantified using the Fiji image processing software (Schindelin et al., 2012). The tissue-specific effect of the EGFP codon usage was analyzed by comparing the normalized EGFP signals in either the AGSC or HPZ between genotypes with a two-tailed unpaired Student’s t-test.

## SUPPLEMENTAL INFORMATION TITLES AND LEGENDS

### Supplemental Figure Legends

**Figure S1. Percentages of genes carrying signatures of natural selection on synonymous codon usage in diverse eukaryotic species, related to Figure 1**. For shown species, all annotated protein-coding genes that passed data pre-processing were included. The percentages of genes carrying selection signatures were corrected by false discovery rate (FDR = 0.05). Species are stacked by phylum and kingdom. Blue, unicellular species; black, multicellular species; red, can be either unicellular or multicellular.

**Figure S2. Clustered codon usage patterns across genes carrying signatures of selection on gene-specific codon usage bias in *A. thaliana*, related to Figure 2.** Hierarchical clustering of all genes identified as under selection for gene-specific codon usage bias in *A. thaliana.* Analyses as in Figure 2.

**Figure S3. Clustered codon usage patterns across genes carrying signatures of selection on gene-specific codon usage bias in S. *cerevisiae*, related to Figure 2.** Hierarchical clustering of all genes identified as under selection for gene-specific codon usage bias in S. *cerevisiae.* Analyses as in Figure 2.

**Figure S4. Clustered codon usage patterns across genes carrying signatures of selection on gene-specific codon usage bias in *C. elegans*, related to figure 2.** Hierarchical clustering of all genes identified as under selection for gene-specific codon usage bias in *C. elegans.* Analyses as in Figure 2.

**Figure S5. Clustered codon usage patterns across genes carrying signatures of selection on gene-specific codon usage bias in *H. sapiens*, related to figure 2.** Hierarchical clustering of all genes identified as under selection for gene-specific codon usage bias in *H. sapiens.* Analyses as in Figure 2.

### Supplemental Table Legend

**Table S1. Gene ontology analysis of gene clusters defined by codon usage patterns, related to Figure 2, Tables 1–2.**

Clusters A and B are defined by their codon usage patterns, and only genes carrying signatures of natural selection on synonymous codon usage are included, as shown in Fig. 2. Molecular function, cellular component, and biological process GO terms are analyzed. Only significant over- or under-representations are shown. Fold enrichment is relative to the entire genome. We did not obtain informative results when analyzing biological process GO terms of Cluster A genes.

**Analytical Solutions of Equation Systems Used to Estimate *μ* Values, Related to Figure 1.**

**Sequences of Fluorescent Reporter cDNAs, Related to Figure 3.**

